# HGCPep: A Hypergraph-based Deep Learning Model for Enhancing Representation of Peptide Features in Cancer-associated ncPEPs Identification

**DOI:** 10.1101/2025.02.13.637760

**Authors:** Wentao Long, Zhongshen Li, Junru Jin, Jianbo Qiao, Yu Wang, Leyi Wei

## Abstract

The emergence of non-coding RNA-encoded small peptides (ncPEPs) has sparked significant interest within the realm of cancer immunotherapy, owing to their potential as valuable therapeutic targets and biomarkers. The identification and characterization of cancer-associated ncPEPs assume a crucial role in propelling cancer research forward and augmenting our comprehension of immune-related processes. However, prevailing methods for cancer-associated ncPEPs identification primarily rely on sequence order, neglecting the latent relationships between peptides. As widely acknowledged in the central dogma, peptides are translated from RNA via a many-to-many mapping. In this study, we capitalize on the translation relationship and introduce a hypergraph-based model called HGCPep to enhance the representation of peptide features. Specifically, RNA is regarded as a hyperedge connecting the peptides that are translated from it. Our experimental results validate that the HGCPep approach, leveraging the strengths of hypergraph and convolutional neural networks, outperforms alternative methodologies in the recognition of ncPEPs. Additionally, employing a reduction tool, we visualize the results of multi-label classification within a 2D latent space, shedding light on how multi-label classification task influence the representation of peptides. The dataset and source code of our proposed method can be found via https://github.com/Longwt123/HGCPep_Github.

## Introduction

Traditionally, non-coding RNAs (ncRNA) were presumed to lack coding potential, and thus earlier investigations focused primarily on their functional roles rather than their coding abilities[1]. However, recent breakthroughs in ncRNAs research have revealed the existence of ncRNA-encoded small peptides (ncPEPs), which have opened up new avenues in cancer research, particularly in the fields of immunotherapy and biomarker identification[2]. Advancements in proteomics technology have uncovered short open reading frames (sORFs) within ncRNAs, thereby elucidating the presence of ncRNA- encoded small peptides (ncPEPs) with significant biological activities[3–6]. This emerging understanding has propelled the exploration of ncRNAs beyond their functional roles, encompassing investigations into their coding potential as well.

Within the realm of cancer research, the discovery of ncPEPs has introduced novel avenues for the identification of therapeutic targets and biomarkers, particularly in the context of cancer immunotherapy[7]. Cancer immunotherapy harnesses the inherent capabilities of the immune system to selectively target and eliminate cancer cells expressing specific antigens. Central to this approach are neoantigens, which arise from genetic alterations in tumor cells and can be recognized by T cell receptors (TCRs), triggering a highly specific anti-tumor immune response[8]. These ncPEPs hold promise as unique neoantigens that could be exploited to develop immunotherapeutic strategies with enhanced specificity and efficacy. Noteworthy examples of ncPEPs as potential biomarkers in cancer have also come to the fore. Chakraborty et al. reported consistent expression of ncPEPs across 11 carcinoma cell lines, suggesting their potential as generalizable biomarkers with remarkable stability[9]. Unraveling the expression patterns and functional implications of ncPEPs in different cancer types holds fundamental importance in deciphering the intricate pathogenesis of cancer and providing crucial insights for precision medicine and targeted therapies.

While protein mass spectrometry (MS[10–12]) remains the primary experimental technique for detecting ncPEPs, computational methods play a crucial role in complementing and enhancing these efforts. However, computational approaches face several challenges that must be overcome to fully exploit the potential of ncPEPs research in cancer. One significant challenge is the accurate association of ncPEPs with specific cancer types, necessitating the development of robust computational frameworks capable of deciphering the intricate relationships between ncPEPs and the molecular landscapes of diverse cancer subtypes. Furthermore, most peptide prediction functions solely rely on the sequence order, such as DeepPep[13] and AlphaPeptDeep[14]. However, they often overlook the inherent information, leading to ineffective and unreliable predictions. In line with the central dogma, which elucidates the translation of peptides from RNA through a many-to-many mapping, the majority of peptides in cancer research are found intracellularly. Leveraging this translation relationship within this context, we can enhance the robustness of our predictions. To harness this additional information, we propose the integration of hypergraphs into our model to better capture this relationship.

In recent years, the utilization of hypergraphs has demonstrated remarkable efficacy in modeling and capturing intricate correlations. The concept of hypergraph learning, initially introduced in [15], can be viewed as a transductive learning process that propagates through the hypergraph structure. This approach has witnessed significant advancements and widespread applications across various domains. In the context of tag-based image retrieval, Wang et al.[16] constructed a sophisticated hypergraph that incorporates global and local visual features alongside tag information to facilitate image relevance learning. This framework leverages the power of hypergraphs to capture complex relationships and enhance the retrieval performance. Building upon the unprecedented achievements of deep learning, researchers have extended hypergraph learning techniques to encompass deep architectures. For instance, Feng et al.[17] introduced hypergraph neural networks (HGNN) to model and learn complex correlations that extend beyond pairwise relationships. In contrast to traditional graph neural networks (GNN[18]), HGNN employs a vertex-hyperedge-vertex information propagation scheme, enabling iterative learning of data representations within the hypergraph structure. The development of deep hypergraph learning techniques exemplifies the fusion of hypergraphs and deep learning, offering enhanced capabilities for capturing intricate correlations and optimizing data representations.

To address the challenges above, we propose HGCPep, an innovative deep-learning framework that combines the power of hypergraphs and convolutional neural networks for the recognition of cancer-associated ncPEPs. By leveraging the unique characteristics of hypergraphs, we construct a hypergraph representation based on the biological relationship between RNA and peptides. Unlike traditional graphs with fixed-degree edges, hypergraphs offer the flexibility to encode complex high-order relationships using degree-free hyperedges. In our framework, each hyperedge represents an RNA molecule, and the connected vertices correspond to its translated peptides. Experimental results on diverse datasets demonstrate the effectiveness and robustness of HGCPep, outperforming other models. Furthermore, our research highlights the embedding enhancement capabilities of hypergraphs, shedding light on cancer pathogenesis and providing valuable insights for cancer research. Additionally, employing a reduction tool, we visualize the multi-label classification results in a 2D latent space, revealing the influence of the multi-label classification task on peptide representation.

## Results

### Hypergraph makes HGCPep prediction more accurate and robust

In order to test the effectiveness of HGCPep on the prediction of multi-label multi-classification problems, we conducted a comprehensive analysis of model prediction AUC scores for each ncPEPs class corresponding to different cancer types, as shown in **Figure 2** below. We used HGCPep as the base model, the prediction AUC of all classes is improved overall after the hypergraph information was incorporated into the model on both two datasets. Being 37.92% higher than the model without hypergraphs, *Thyroid cancer* class exhibits the most noticeable improvement in the 15 classes dataset. In the 10 classes dataset, *Leukemia* class significantly outperformed the model without hypergraph, growing by 22.88%. These results show that hypergraph structure and hypergraph convolution module play a critical role in improving the prediction accuracy of each class in this problem.

**Figure 1.**
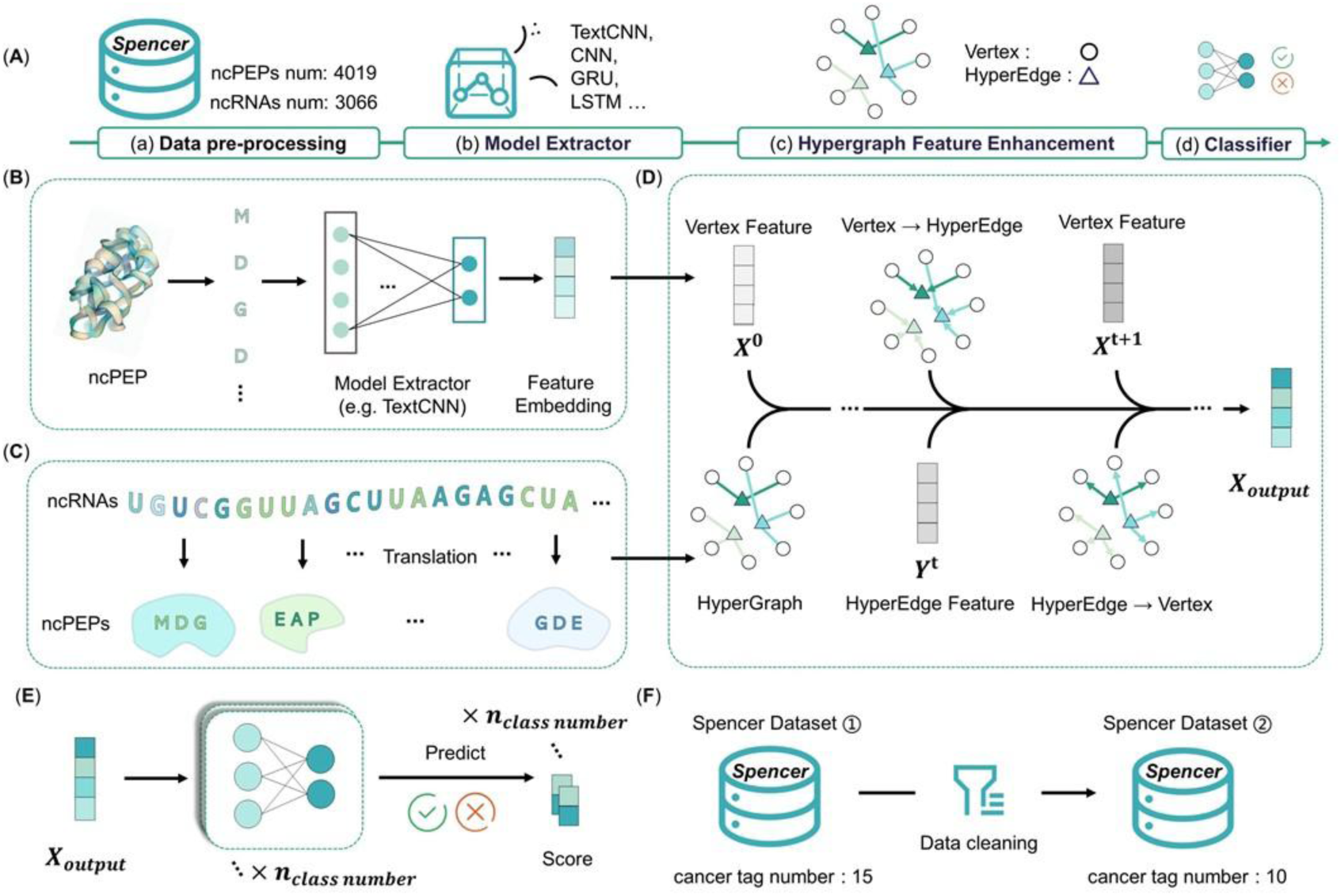
The framework of the HGCPep method for ncPEPs prediction. **A.** The overall workflow schema of the HGCPep method for ncPEPs prediction, which composes of four blocks: (a) data pre-processing block, (b) model extractor block, (c) hypergraph feature enhancement block, and (d) classifier block. The model extractor block extracts features as shown in (**B**), while the hypergraph feature enhancement block constructs hypergraph and strengthens feature representation according to (**C**) and (**D**) respectively. In (**D**), the circles represent hypergraph nodes, the triangles represent hyperedges, x represents the features of nodes, and y represents the features of hyperedges. (**E**) is the classifier we use and (**F**) is the two datasets we build.

**Figure 2.**
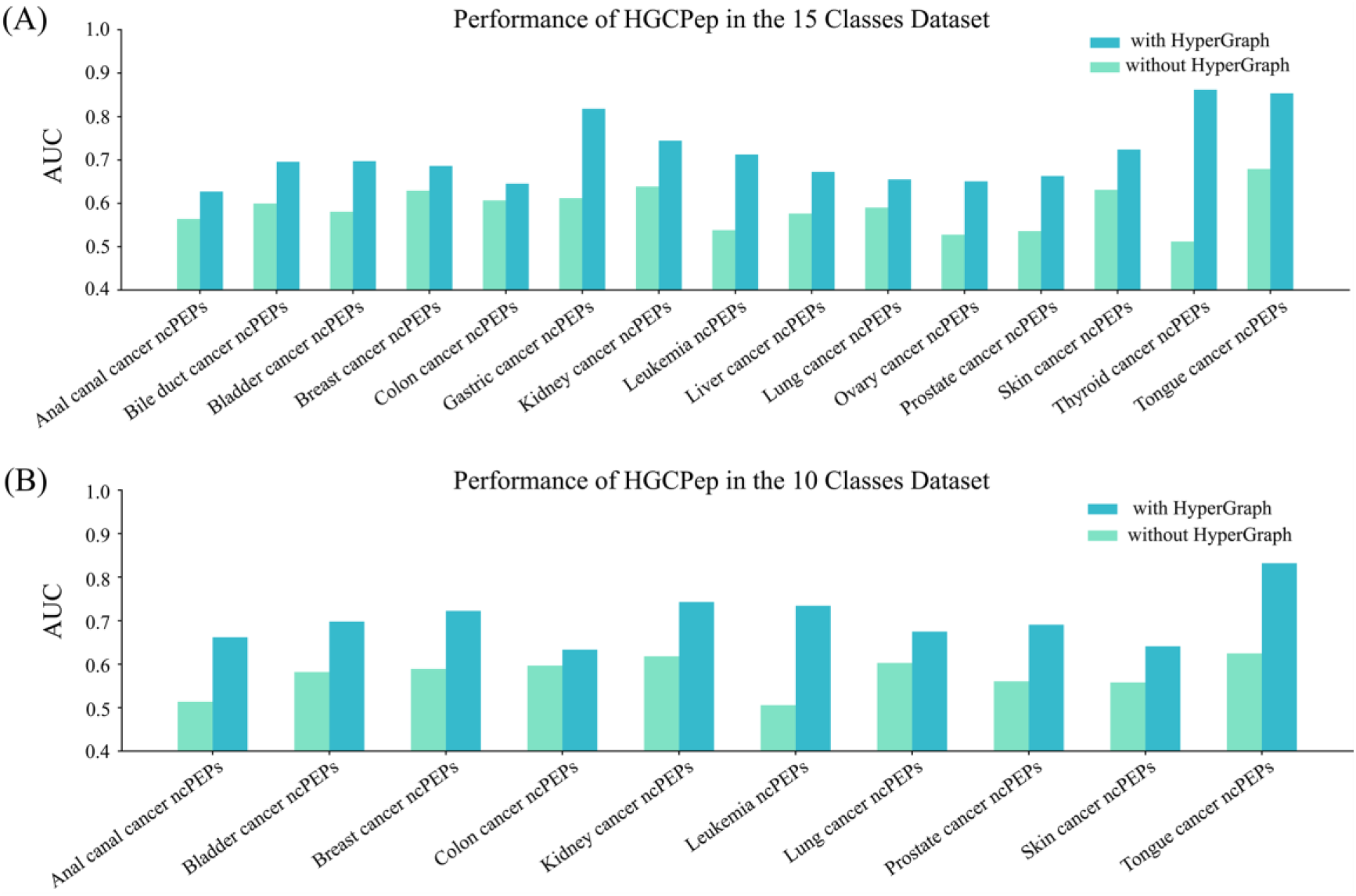
Performance evaluation on the accuracy of HGCPep. Performance evaluation on the accuracy of HGCPep for predicting ncPEPs in various types of cancers in (**A**) the 15 classes dataset and (**B**) the 10 classes dataset.

For demonstrating that the enhancement capability of hypergraph is not limited to our model, we combine hypergraph convolution with other commonly used embedding models, and compare the prediction results. Here, we choose some popular methods, including CNN[19], GRU (Gate Recurrent Unit[20]), LSTM (Long Short-Term Memory[21]), LSTM with Attention[22], RNN and CNN (Recurrent neural network[23]). Through the comparison of MCC, ACC and AUC metrics in **Figure 3**, we find that the hypergraph information is applied to improve the feature embedding of the other models above, and this model can also achieve excellent performance improvements. The results show that hypergraphs can significantly improve the performance of various models, as shown in Figure 3 (A) has an average increase of 9.72%, 23.16% and 12.22% in MCC, ACC and AUC in the 15 classes dataset, and (B) has an average increase of 6.65%, 20.91% and 12.64% in MCC, ACC and AUC in the 10 classes dataset, respectively. Models without hypergraphs exhibit high ACC scores, but their Precision scores are notably low. These results indicate that hypergraphs can obtain the high-dimensional potential characteristics of ncPEPs in corresponding ncRNAs, thus improving the expression ability of feature embedding.

**Figure 3.**
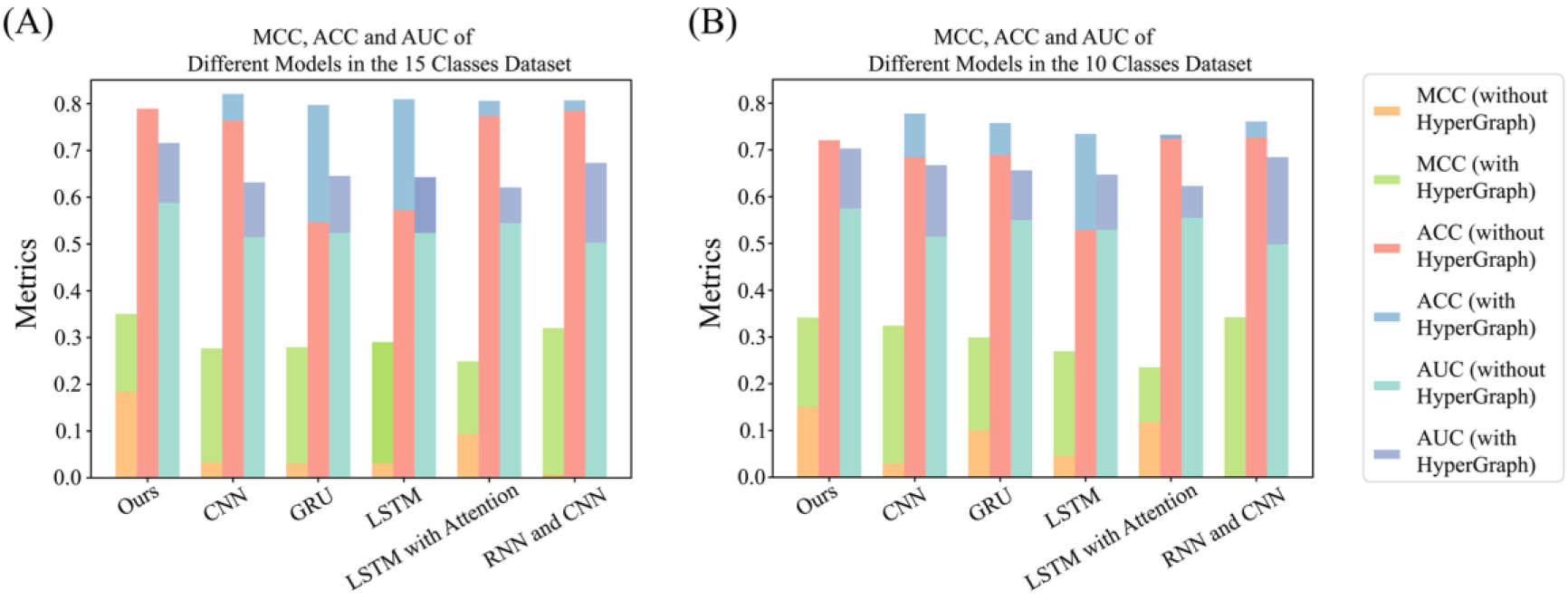
Performance evaluation on the MCC, ACC and AUC of HGCPep. Performance evaluation on the MCC, ACC and AUC of HGCPep, CNN, GRU, LSTM, LSTM with Attention, RNN and CNN for predicting ncPEPs in (**A**) the 15 classes dataset and (**B**) the 10 classes dataset.

### Performance comparison of different hypergraph convolution models

To compare the influence of different types of hypergraph convolution on prediction results, we employ TextCNN as the basic model, and conducted comparative experiments with HGNN hypergraph convolution model and HGNNP hypergraph convolution model according to the study of Gao et al.[24]. The comparison results of AUC are shown in **Figure 4** below. From these results we can have the following observations: (1) Hypergraphs significantly improve the model prediction performance. The average performance improvement of the two hypergraph convolution models can reach 12.80% and 11.56% in the 15 classes dataset. The increase reached its peak at an astonishing 37.92%. Improvements are also made in the 10 classes dataset. (2) In comparison to HGNN and HGNNP hypergraph convolutions, HGNNP demonstrates improved performance across 6 classes, similar performance in 7 classes (with differences not exceeding ±2.0%), and decreased performance in 2 classes within the 15 classes dataset. For 10 classes dataset, the corresponding numbers are 2, 6, and 2 classes, respectively. In conclusion, the HGNNP hypergraph convolution module is more effective for the prediction of ncPEPs of cancer compared to the HGNN module. **Performance comparison of different graph convolution models**

**Figure 4.**
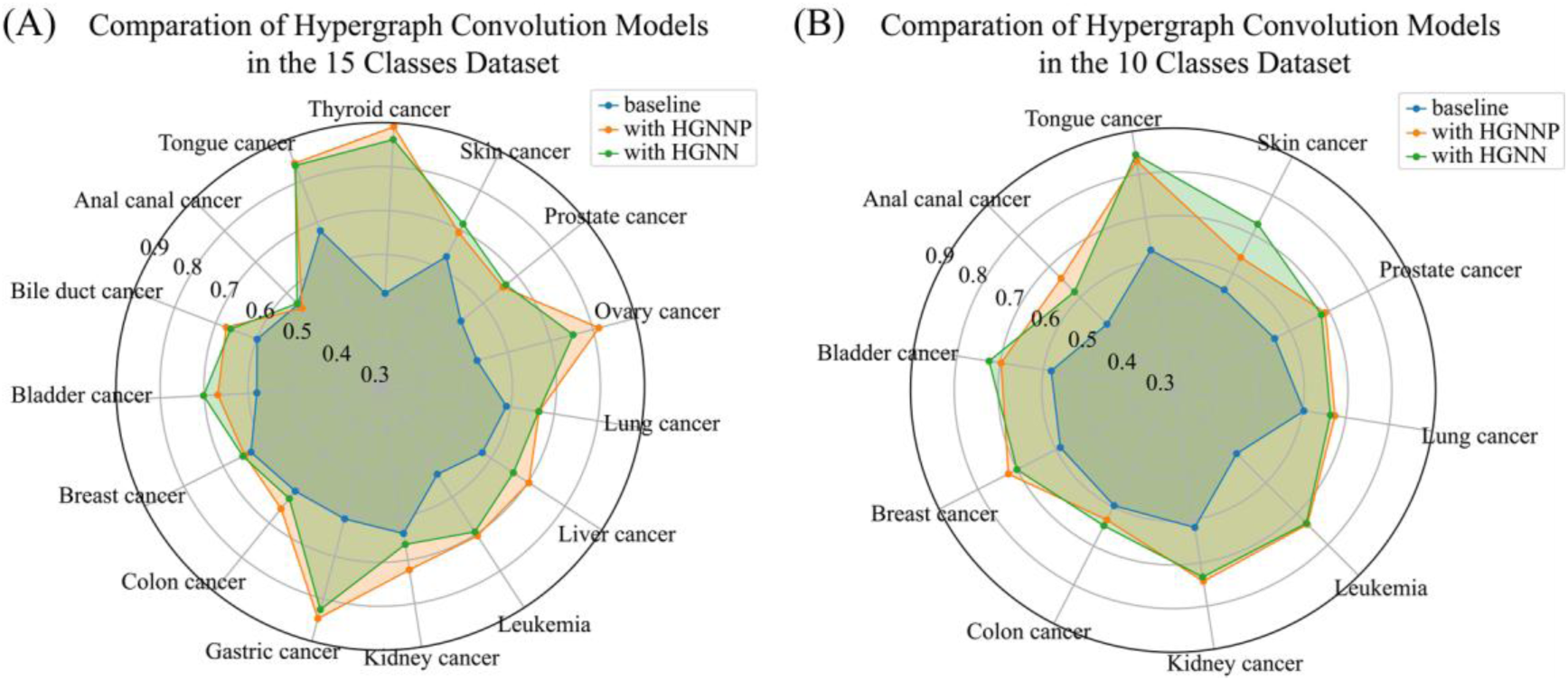
Performance evaluation on the AUC of the various hypergraph modules. Performance evaluation on the AUC of the various hypergraph modules employed by HGCPep for forecasting ncPEPs in various cancers in (**A**) the 15 classes dataset and (**B**) the 10 classes dataset.

To compare the impact of different types of graph convolutions on the prediction results, we used TextCNN as the base model and conducted comparative experiments with classic graph neural network models based on the study by Huang et al.[25]. As **Figure 5** (A) and (B) illustrated, our hypergraph model outperformed the classic ordinary graph models. Additionally, we investigated potential overfitting of the model by comparing models with different dropout parameters, confirming that our model did not exhibit significant overfitting in Figure 5 (C).

**Figure 5.**
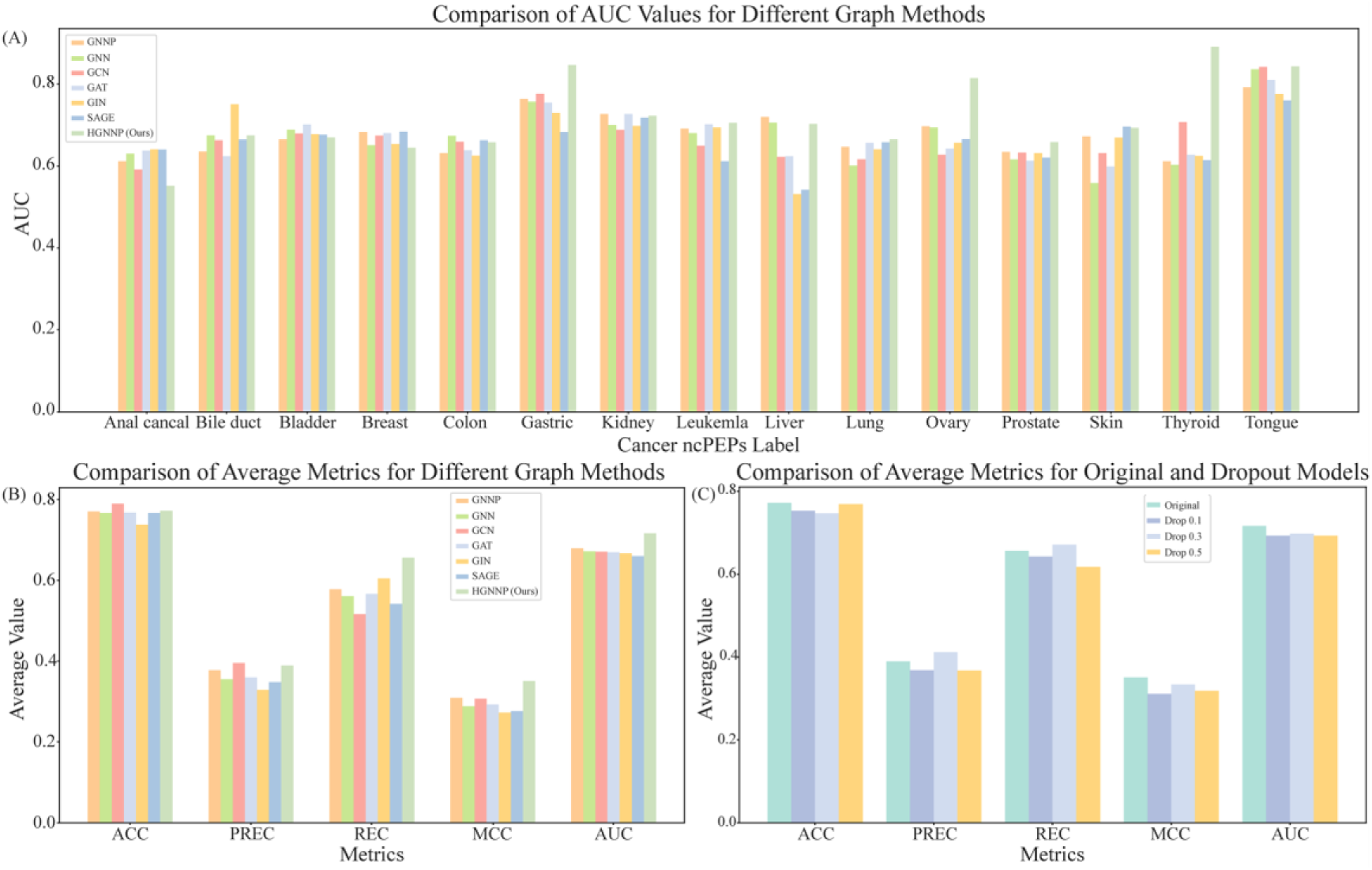
Performance evaluation of the various graph modules for forecasting ncPEPs. Performance evaluation of the various graph modules for forecasting ncPEPs in various cancers in (**A**) and (**B**). Performance evaluation of different dropout parameters in (**C**).

### Performance comparison between different loss functions

To overcome the imbalance in multi-label classification of ncPEPs, we employ several special loss functions in the HGCPep model’s training process. Here, we compared the performance of different loss functions to optimize HGCPep, including OHEM (Online Hard Example Mining[26]), commonly used Focal Loss, Cross Entropy Loss (CE), and CE with weight. By recording and comparing the AUC indexes of different loss in the training process, we compared the performance of HGCPep under different loss functions, as shown in **Figure 6**. To be noticed, the line has been processed by the gaussian smooth. As can be seen from Figure 6 (A) and (B), the training process of our model on the two datasets is stable without large fluctuations, and the overall effect is good. The horizontal axis in Figure 6 (A) represents the maximum values of different loss types, illustrating the exceptional performance of the OHEM loss in the 15 classes dataset. With the dataset being cleaned to reduce its imbalance, the prominence of the OHEM loss diminishes in the 10 classes dataset, as depicted in Figure 6 (B). To better learn which loss function work in the two datasets, we compare different combination of three loss function in the Figure 6 (C) and Figure 6 (D). The impact of the OHEM technique on the imbalanced dataset is more pronounced in the 15 classes dataset compared to the 10 classes dataset. In the imbalanced dataset, various loss functions exhibit unstable performance, making it challenging to determine the optimal choice. However, in the 10 classes dataset, all specialized loss functions demonstrate improvement over the original CE loss, indicating the effectiveness of this dataset for deep learning training applications.

**Figure 6.**
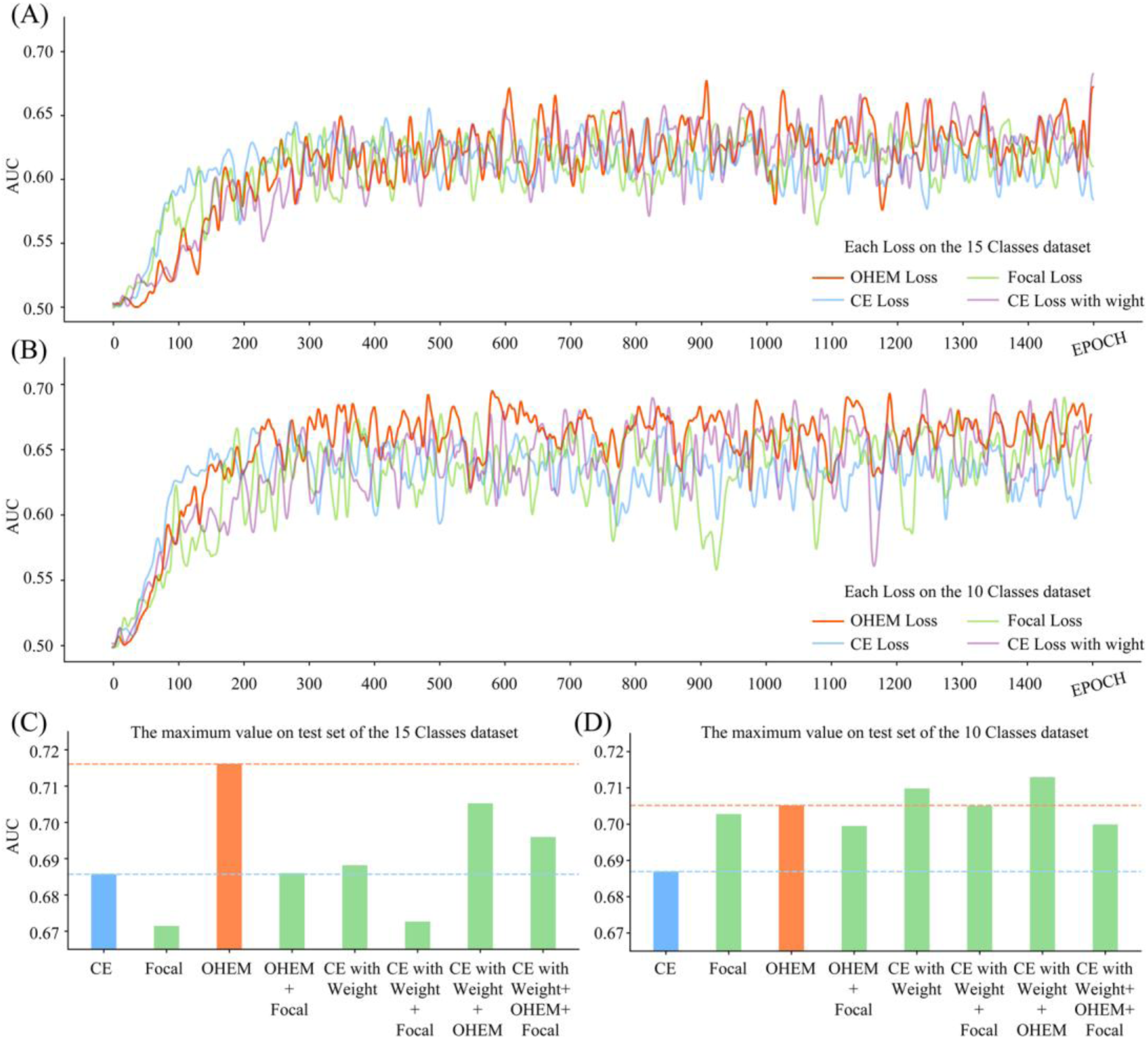
Performance evaluation on the AUC of HGCPep. Performance evaluation on the AUC of HGCPep for predicting ncPEPs in various types of cancer in (**A**)(**C**) the 15 classes dataset and (**B**)(**D**) the 10 classes dataset.

### Visualization of HGCPep

Considering that both datasets naturally have hypergraph structure, which indicates that they contain complex high-order correlations between ncPEPs data. In order to visually demonstrate the learning capability of the hypergraph-based methods, we visualized the optimal model of HGCPep. The output of the model was visualized in Euclidean Space[24], and the results were shown in **Figure 7**. Among them, different colors represent different multi-label categories of data, and their dimensions are reduced to two dimensions for the distribution of (A) and (B) in Figure 7. The multi-label categories represented by (C) in Figure 7 correspond to (A). Each row of Figure 7 (C) is one class in the 15 classes dataset, and each column is one of the multi-label categories, which contains one or more points representing multiple labels. The categories represented by the colors in Figure 7 (A) correspond one-to-one with the categories represented by the colors in Figure 7 (C). In addition, the number of each category is illustrated in bar graphs in Figure 7 (C). Similarly, Figure 7 (D) corresponds to Figure 7 (B). It can be seen from the results that our hypergraph-based method produces discernible clustering, qualitatively verifying the effectiveness of the proposed method.

**Figure 7.**
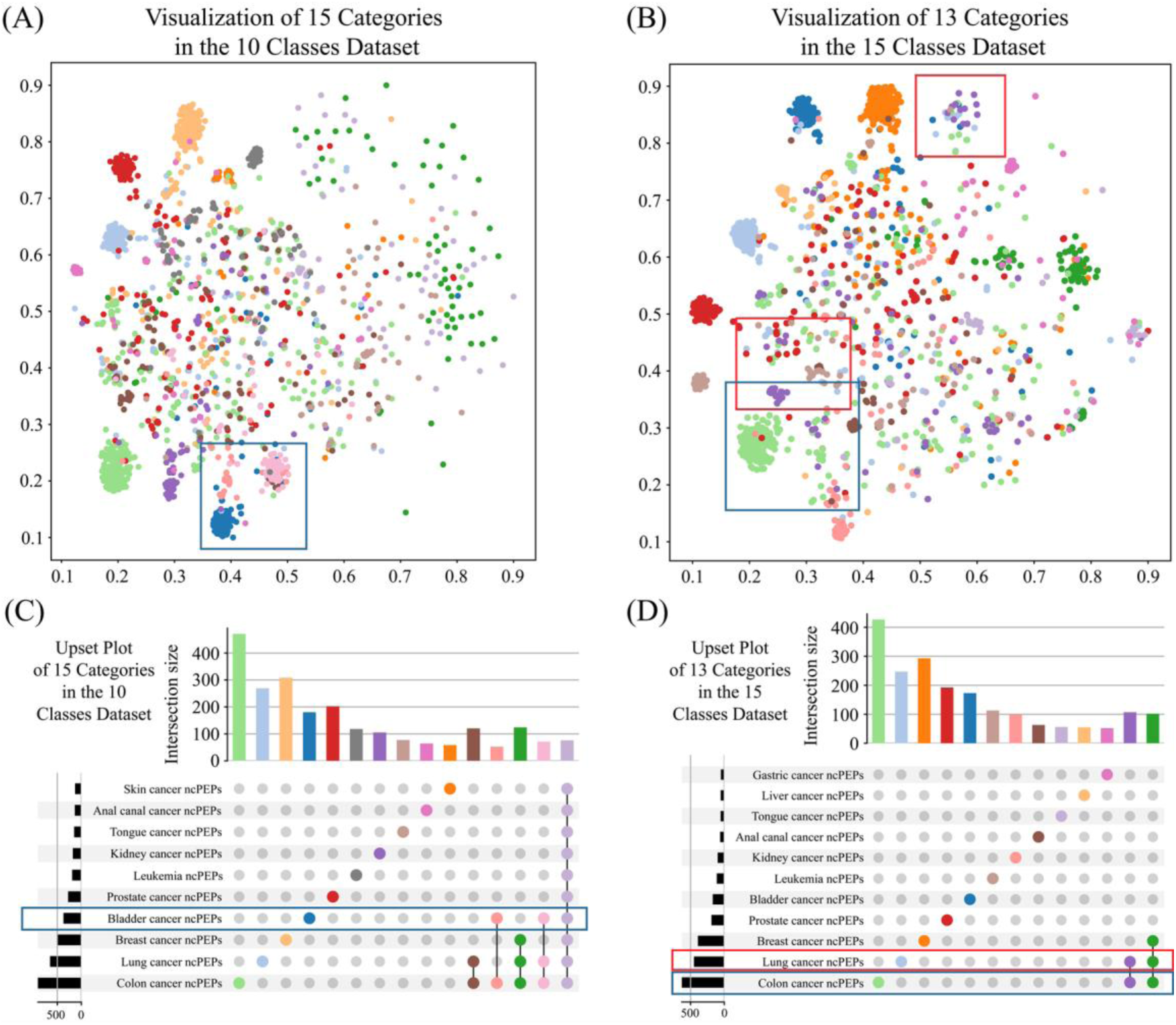
Evaluating HGCPep in euclidean space. Evaluating HGCPep in euclidean space in (**A**)(**C**) the 10 classes dataset and (**B**)(**D**) the 15 classes dataset. In (**A**) and (**C**), the different colored dots and bars represent cancer categories, and the same applies to (**B**) and (**D**). In the tables shown in (**C**) and (**D**), each row represents a cancer label, with the bar on the left indicating the number of instances for that label; Each column represents a multi-label combination, with the bar on top indicating the number of instances for that combination.

About visualization results in the 10 classes dataset, in the highlighted region within blue box of Figure 7 (A), data points labeled in blue, light pink, and pink tend to cluster together, suggesting that they share a common label, *Bladder cancer* which is further supported by the ninth row in Figure 7 (C). As for the dark green and light purple labeled data points that do not exhibit clear clustering patterns in the figure, we speculate that their difficulty in achieving visible clustering results stems from their involvement with three different label categories and their similarity to many other labeled data points. Consequently, the model faces challenges in accurately classifying them, leading to less distinct visual clustering effects.

About visualization results in the 15 classes dataset, the partial region highlighted in red box in Figure 7 (B) demonstrates the clustering results of the purple-labeled data and light blue-labeled data, which exhibit a close resemblance. Through careful analysis and statistical examination, as indicated in the ninth row of Figure 7 (D), these data points share a common label, *Ovary cancer*, resulting in similar clustering outcomes after dimensionality reduction. Similarly, in blue box, the purple-labeled data and light green-labeled data exhibit proximity in their clustering results, sharing the common label, *Colon cancer*. In addition, We have considered and conducted other visualization methods[27] and comparative experiments[28–30], as detailed in the appendix.

## Conclusion

In this study, we present HGCPep, an innovative deep-learning framework that combines the power of hypergraphs and convolutional neural networks for the recognition of cancer-associated ncPEPs. By leveraging the unique characteristics of hypergraphs, we construct a hypergraph representation based on the biological relationship between RNA and peptides. Unlike traditional graphs with fixed-degree edges, hypergraphs allow for encoding complex high-order relationships using degree- free hyperedges. Our experimental results on diverse datasets demonstrate the effectiveness and robustness of HGCPep, surpassing other existing models. Furthermore, we highlight the embedding enhancement capabilities of hypergraphs, providing valuable insights into cancer pathogenesis and contributing to cancer research.

To address data imbalance, we organize our dataset based on commonly used databases in the field and construct a cleaned dataset according to **Table 1**, mitigating the impact of data imbalance. Comparative experiments, as shown in Figure 4, confirm that the HGNNP hypergraph module is particularly suited to address the current problem. Additionally, we introduce the OHEM loss function to alleviate data imbalance, and the results in the line chart of Figure 5 demonstrate its effectiveness. Notably, as the dataset becomes slightly more balanced, the improvement from the OHEM loss function becomes less pronounced. Furthermore, by employing a reduction tool, we visualize the multi-label classification results in a 2D latent space, offering insights into the influence of the multi-label classification task on peptide representation. All in all, HGCPep can be an important tool to discover cancer-associated ncPEPs and give ideas on how to make use of intracellular information.

**Table 1.** Information about the two datasets.

|  | 15 Classes Dataset | 10 Classes Dataset |
| --- | --- | --- |
| *Anal canal cancer* | 518 | 518 |
| *Bile duct cancer* | 442 | - |
| *Bladder cancer* | 1178 | 1178 |
| *Breast cancer* | 1075 | 1075 |
| *Colon cancer* | 2068 | 2068 |
| *Gastric cancer* | 460 | - |
| *Kidney cancer* | 905 | 905 |
| *Leukemia* | 506 | 506 |
| *Liver cancer* | 244 | - |
| *Lung cancer* | 1646 | 1646 |
| *Ovary cancer* | 258 | - |
| *Prostate cancer* | 1028 | 1028 |
| *Skin cancer* | 552 | 552 |
| *Thyroid cancer* | 200 | - |
| *Tongue cancer* | 519 | 519 |

Note: There were 4,019 peptides, 3,066 RNAs (both), and 11,599 labels (about the 15 classes dataset). ‘-’ indicates that it has been removed.

## Materials and methods

### Dataset

We construct the cancer-associated ncPEPs dataset from the SPENCER database[31]. Currently, SPENCER has collected a total of 2806 mass spectrometry (MS[10–12]) data points from 55 studies, encompassing 1007 tumor samples and 719 normal samples. Through the utilization of an MS-based proteomic analysis pipeline, SPENCER has identified 29,526 ncPEPs in 15 different cancer types, with 22,060 of these ncPEPs experimentally validated in other studies. This comprehensive dataset serves as a commonly used and valuable resource for ncPEPs-related investigations. We classified and curated all samples, resulting in a dataset comprising 4019 samples, each containing an amino acid sequence and a label list indicating the associated cancer types. In consideration of the association of a single peptide with multiple cancer types, we designed a dataset comprising a multi-label classification task. However, the presence of imbalanced distribution among certain cancer types within the dataset poses challenges to effective model training. To address this issue, we implemented a filtering process, excluding cancer tags with a count lower than 500, thereby constructing a secondary dataset (as illustrated in Table 1 and **Figure 1** (F)).

#### Hypergraph Building and Data Pre-Processing

To build the hypergraph, we treat peptides as nodes and connect them using hyperedges that correspond to multiple peptides translated by the same RNA. This design also allows for a peptide to be potentially translated from multiple RNAs, meaning a node could be associated with multiple hyperedges. This structure captures the complex relationships between peptides and RNAs, reflecting the multiple translations and interactions within the biological system.

The original peptide data included the serial number, sequence information and corresponding cancer label information of ncPEPs. First, it is mapped into *psDict* = {*pid*: *sequence*} and *plDict* = {*pid*: *labels*} dictionaries, where *pid* is the sequence number of ncPEPs, *sequence* is its sequence information, and *labels* is its label information. The original RNA data included the sequence information of ncPEPs and the number of corresponding RNA. We counted the set of RNA numbers *rDict* = {*rid*: *frequency*} , where *rid* was the sequence number of RNA and frequency denotes how often each RNA appears in the original RNA data, and mapped the data into *cDict* = {*pid*: *edges*} using *psDict* , where *edges* = [*rid*1, *rid*2. . . ] are multiple RNAs corresponding to *pid*.

Next, we map each ncPEPs to the vertices of the hypergraph while recording the number of vertices *num*_*vertices*_ and generating corresponding tags through *plDict*. We constructed each RNA hyperedges by *rDict* and *cDict* , concatenating information about corresponding vertices. Finally, a complete Hypergraph was constructed by ***G*** = *Hypergraph*(*num*_*vertices*_, *edges*) (where *Hypergraph* is the function for constructing the hypergraph and ***G*** is the hypergraph we constructed[17]), which is ready for the downstream tasks.

#### Train/valid/test Split

For each of our two datasets, we performed a three-way segmentation, dividing them into distinct sets. This process allocated 80% of the samples for training, 10% for validation, and the remaining 10% for testing. Following the validation phase, we meticulously selected the best-performing model based on the evaluation metrics obtained during this stage. Subsequently, the chosen model underwent thorough testing on the independent test set to assess its performance and acquire reliable metrics.

### Framework Overview

Within this section, we offer a brief overview of our framework HGCPep. HGCPep is designed for representation learning and predicting the type of cancer associated with ncPEPs using raw ncPEPs sequences and corresponding RNA data. The flow chart of our framework is shown in Figure 1 (A), which includes four steps: (a) data pre- processing block, (b) model extractor block TextCNN (Text Convolution Neural Network)[32], (c) hypergraph feature enhancement block[17, 24], and (d) classifier block. For peptide sequence data, the TextCNN module is first used to facilitate feature extraction, extracting key features of the peptide sequence, as shown in Figure 1 (B). Subsequently, the innovative introduction of the HyperGraph structure further enhanced the exploration of the potential relationship between peptides and RNA, where the construction of hypergraph and the application of hypergraph convolution are shown in (C) and (D) of Figure 1. The hypergraph convolution[24] is then applied to elevate the correlation features to generate a more robust representation of our problem. Finally, like (E) in Figure 1, the latent features are obtained through the fusion of key features and association features of peptide sequences, and then utilized by a multi-label multi-classification classifier to predict the corresponding cancer types. In addition, Figure 1 (F) are the two data sets we used, which contain data from 15 and 10 classes respectively.

#### HGNNP module

In this section, we introduce hypergraph structure into our model through RNAs that transcribe peptides to solve the problem of association information between ncPEPs. In an ordinary graph, each edge is limited to connecting only two vertices. However, hypergraphs offer a higher degree of flexibility and expressiveness as they allow hyperedges to connect more than two vertices. This feature allows for the creation of various types of hyperedges when dealing with data that exhibits many-to-many relationships. Consequently, the hyperedges can be seamlessly integrated into a hypergraph, providing an efficient and effective means of data representation. This approach allows for a more nuanced and sophisticated understanding of the complex interactions that characterize many-to-many relationships, and can enable researchers and analysts to uncover new insights and discoveries in a wide range of fields.

After deeply understanding the concept of hypergraph, we have concluded that the existing data is inherently linked to the graph structure, which is very suitable for the application direction of hypergraph. First, we collated the data set, recorded the number of peptides as the number of hypergraph vertices. We then extracted the available paired data correlations and constructed the hyperedge data accordingly. Here we let **G**_*s*_ = (**V**_*s*_, **E**_*s*_) indicate the hypergraph structure, where *v*_*i*_ ∈ **V**_*s*_ is a vertex and 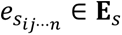 is a hyperedge in the hypergraph connecting *v*_*i*_, *v*_*j*_, ⋯ , *v*_*n*_. Where 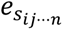 is denoted by 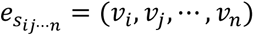 and **E**_*s*_ is expressed as:

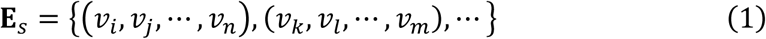

Let *N* denotes the number of hypergraph vertices. Given such *N* and **E**_*s*_ , the hypergraph **G**_*s*_can be generated as follows:

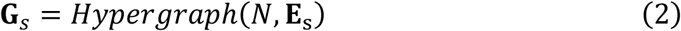

Next, Gao et al. proposed the hypergraph convolution model HGNNConv+[24]. According to the DHG toolkit documentation, we first establish a general hypergraph convolution layer known as HGNNP. This innovative layer comprises a two-phase messaging framework that contains two-phase aggregation functions. In HGNNP, there are two model lists named HGNNPConv, each containing fully connected layers, normalization layers, feature update layers, and dropout layers. The feature update layers include an aggregation-update layer from nodes to edges and an aggregation- update layer from edges to nodes, similar to GNNs. These structures enable HGNNP to be more flexible and perform well in downstream tasks.

The details of HGNNP utilized in our model can be observed in the following formula. Where **X**^t^ represents the feature set of input vertices at layer *t*. The transpose of **H** empowers the control of each vertex feature in the **X**^t^ with respect to its hyperedge neighbors. Therefore, it can be used to guide the aggregation of each vertex, ultimately producing the hyperedge feature set **Y**^t^, which can be represented mathematically as 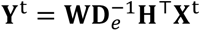. The updating of the vertex feature set **X**^*t*+1^ from the hyperedge feature set **Y**^t^ can be expressed as 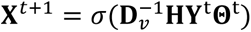. Therefore, the matrix representation of HGNNP can be denoted as:

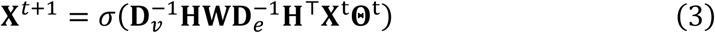

#### Text Convolution Neural Network

The Text Convolution Neural Network (TextCNN[32]) shares similarities with the Convolution Neural Network (CNN[19]) structure. It is a modified version of CNN that leverages CNN’s parallel computing capabilities to accelerate training speed. In addition to preserving the original characteristics of CNN, TextCNN also incorporates text feature extraction, making it applicable for extracting peptide sequence features in our study. The resemblance between text and peptide sequence allows TextCNN to effectively identify linguistic n-grams in the task. Moreover, its convolution structure enables the sharing of predicted behavior among n-grams that have similar elements, even if a particular n-gram has not been encountered before during prediction. Consequently, the multi-channel convolution mechanism in TextCNN automatically captures both local and global information from the sequence, facilitating the learning of peptide sequence features that are specific to our task.

### Performance Measures

Following the previous works, we use following classification metrics to evaluate our model: accuracy (*ACC*), Matthew correlation coefficient (*MCC*) and *AUC* (Area Under Curve of Receiver Operating Characteristic). The formulas of these metrics are described as follows:

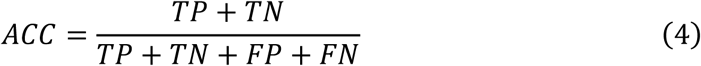

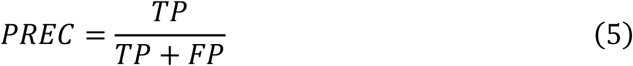

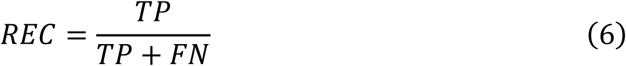

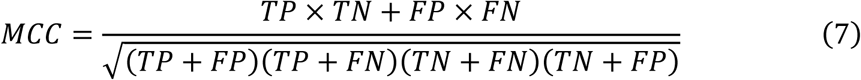

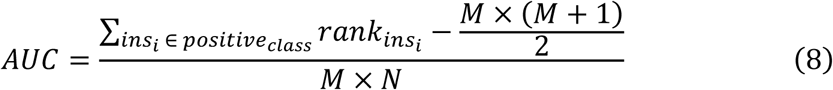

In the formulas, *TP* is the number of true-positive samples, *FP* is the number of false-positive samples, *TN* is the number of true-negative samples and *FN* is the number of false-negative samples. *ACC* is the ratio of the correctly classified samples to the total samples, which is the most intuitive performance measurement. Precision (*PREC*) shows the accuracy of the positive predictions, and Recall (*REC*) measures how well the model identifies all the positive samples. *MCC* takes into account positives and negatives, so that it is generally regarded as a balanced metric to measure the overall performance of a model. In the formulas, 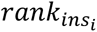 represents the order from smallest to largest of the probability scores for the sample *i* , and 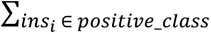 stands for summing only the serial numbers of the positive sample.

*M* and *N* are the number of positive and negative samples respectively. Another metric called the area under the receiver operating characteristic curve (*AUC*) is used for evaluation. *AUC* represents the probability that a random positive is positioned to the right of a random negative, calculated from the area under the receiver operating characteristic curve (ROC), and generally ranges from 0.5 to 1.

## Authors’ contributions

LWT, LZS and JJR conducted ncPEP feature research, participated in the design and construction of the study, conducted statistical analysis, and drafted the manuscript. Others participated in the design and coordination, and helped draft the manuscript. All authors have read and approved the final manuscript.

## Acknowledgments

The work was supported by the Natural Science Foundation of China (Nos. 62071278).

---

**Figure S1.**
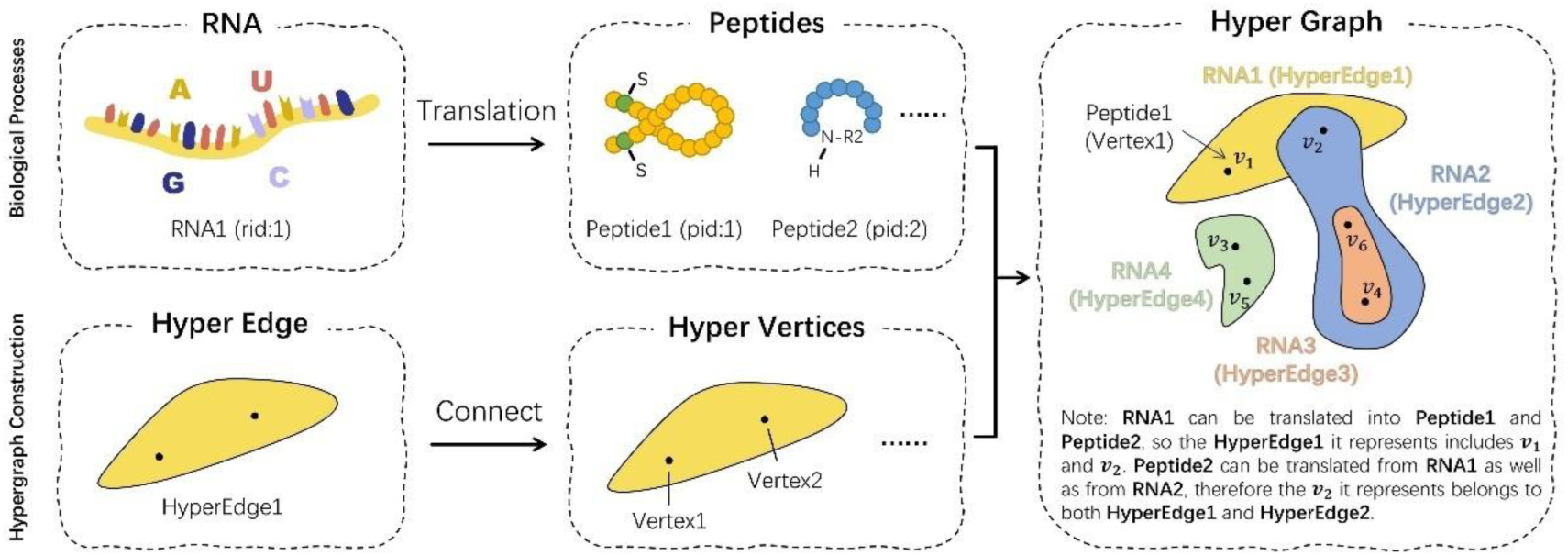
Example of hypergraph construction on the dataset.

**Figure S2.**
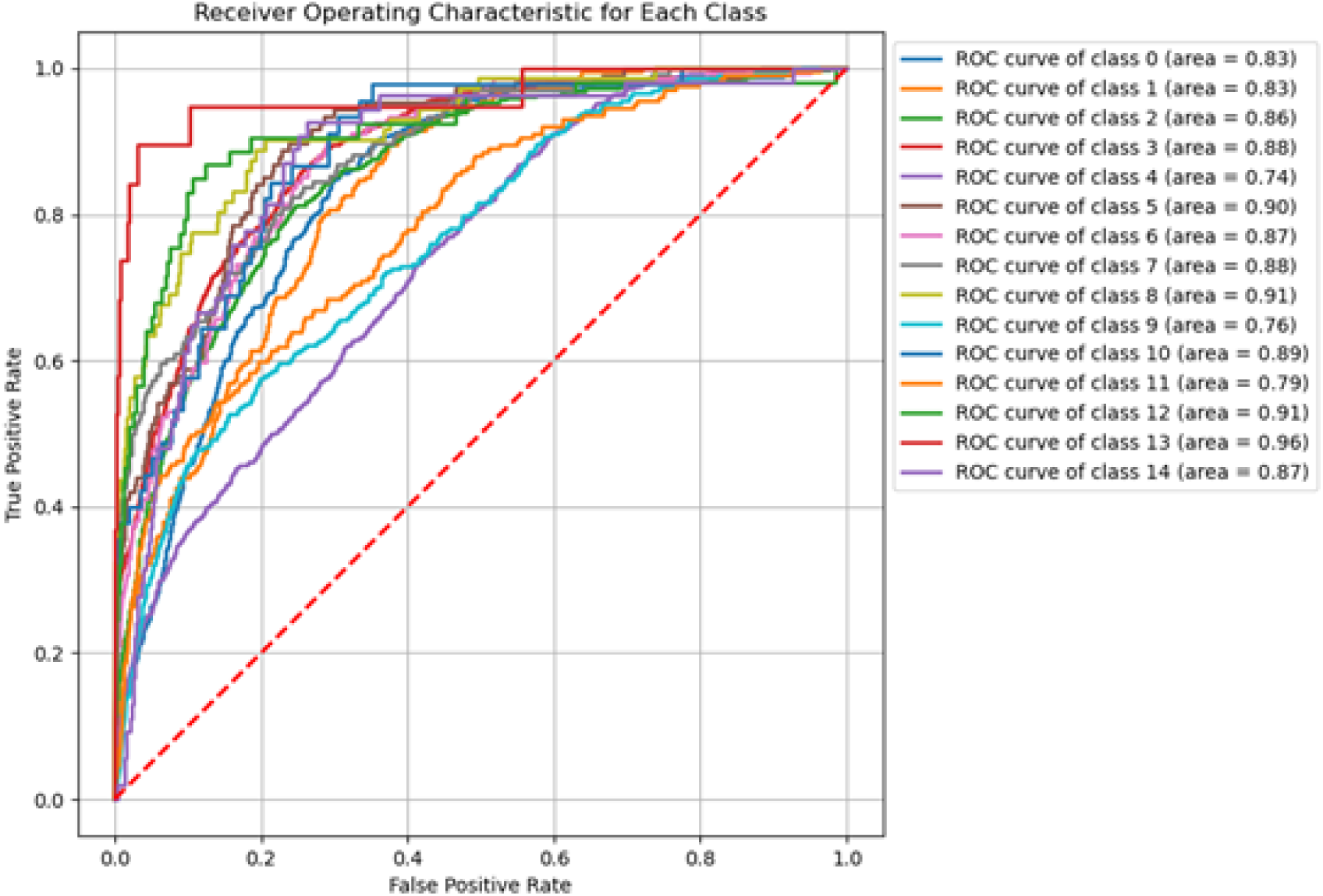
Receiver operating characteristic for each class of HGCPep in the 15 classes dataset.

**Figure S3.**
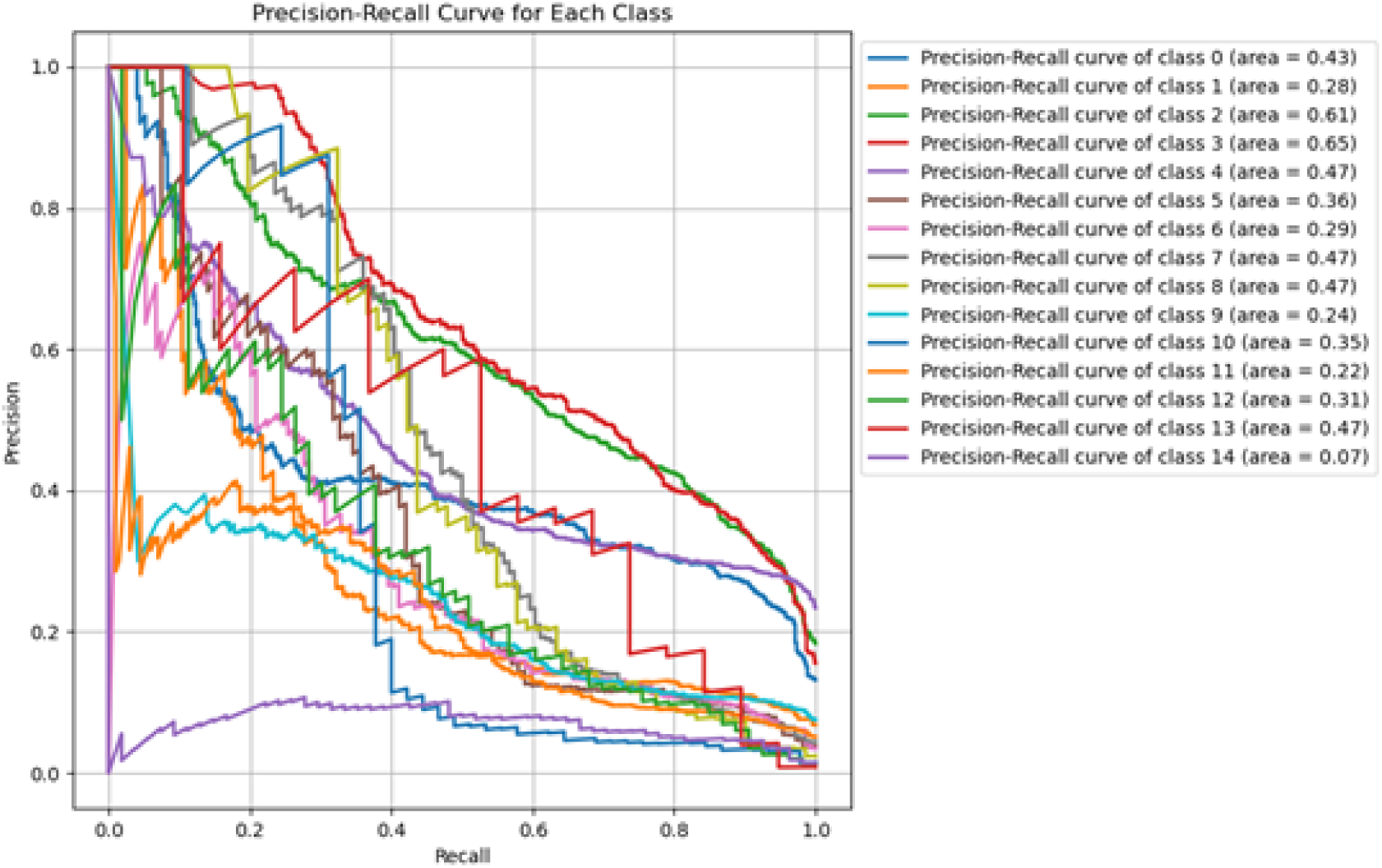
Precision-Recall curve for each class of HGCPep in the 15 classes dataset.

**Figure S4.**
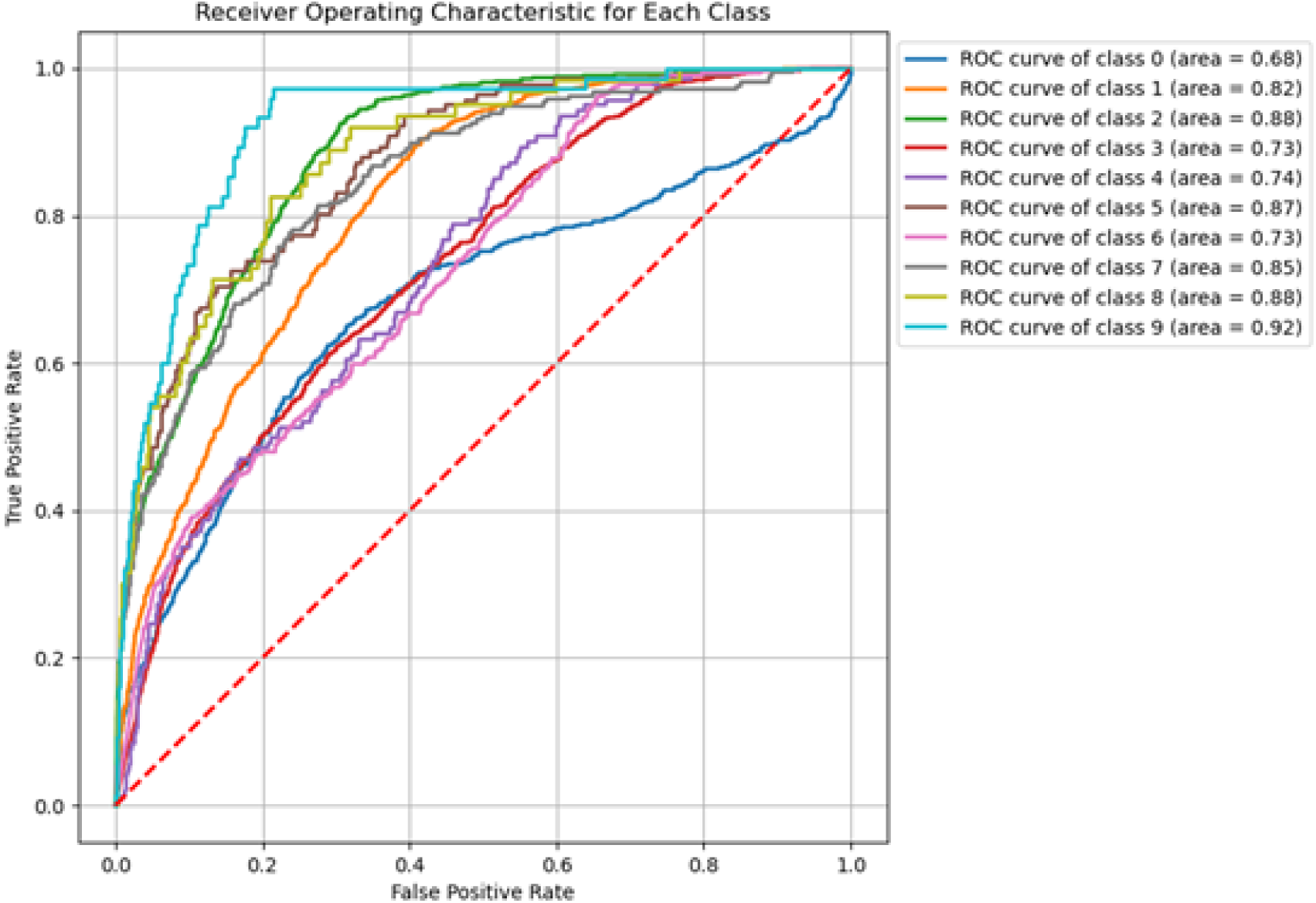
Receiver operating characteristic for each class of HGCPep in the 10 classes dataset.

**Figure S5.**
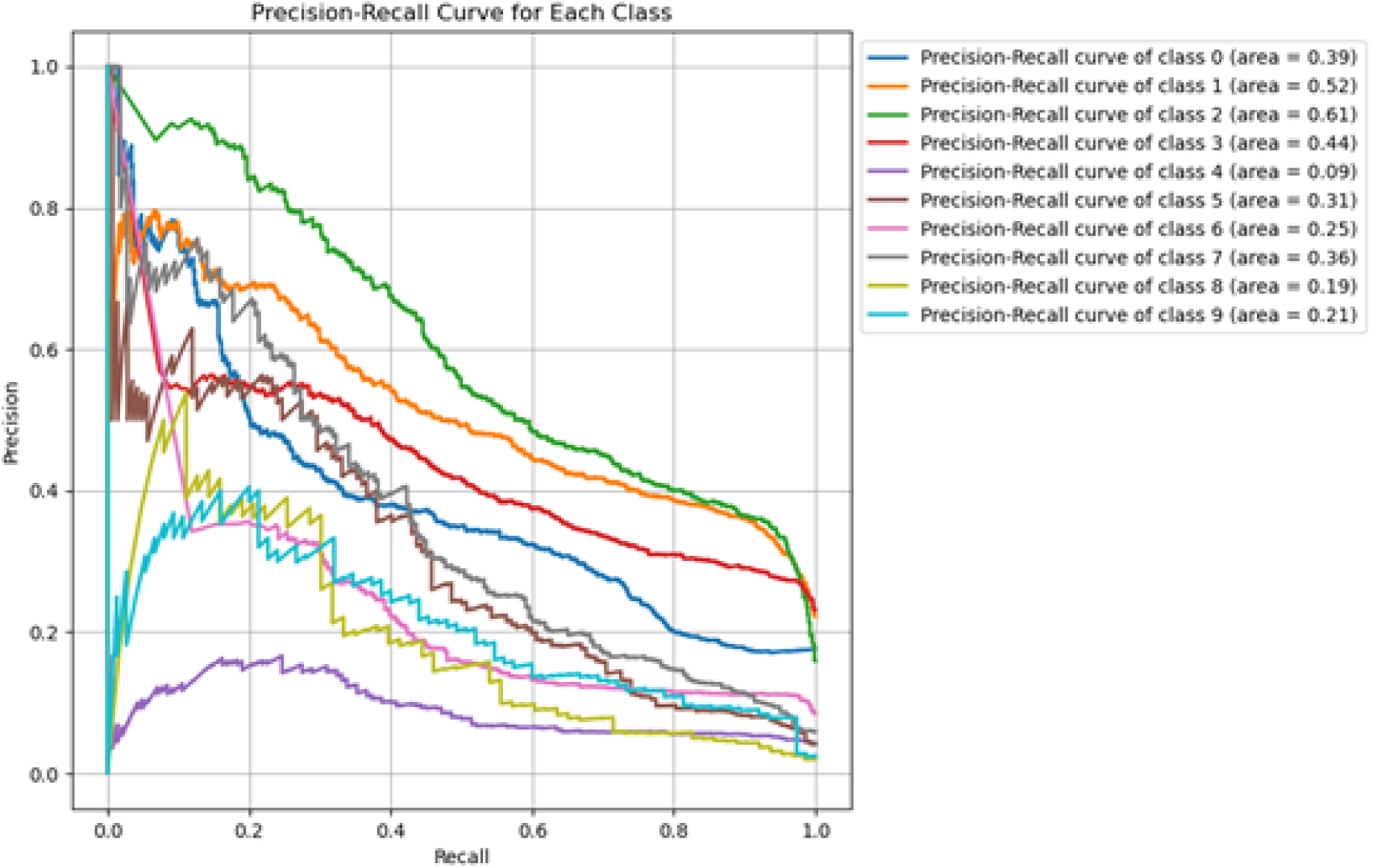
Precision-Recall curve for each class of HGCPep in the 10 classes dataset.

**Table S1.** Performance evaluation on the accuracy of HGCPep for predicting ncPEPs in various types of cancers in the 15 classes dataset.

|  | without HyperGraph | with HyperGraph |
| --- | --- | --- |
| *Anal canal cancer* | **0.5639** | **0.5517** |
| *Bile duct cancer* | **0.5995** | **0.6744** |
| *Bladder cancer* | **0.5806** | **0.6695** |
| *Breast cancer* | **0.6293** | **0.6452** |
| *Colon cancer* | **0.6066** | **0.6579** |
| *Gastric cancer* | **0.6119** | **0.8466** |
| *Kidney cancer* | **0.6387** | **0.7225** |
| *Leukemia* | **0.5382** | **0.7054** |
| *Liver cancer* | **0.5764** | **0.7027** |
| *Lung cancer* | **0.5902** | **0.6653** |
| *Ovary cancer* | **0.5277** | **0.8142** |
| *Prostate cancer* | **0.5360** | **0.6586** |
| *Skin cancer* | **0.6311** | **0.6927** |
| *Thyroid cancer* | **0.5119** | **0.8911** |
| *Tongue cancer* | **0.6793** | **0.8434** |

**Table S2.** Performance evaluation on the accuracy of HGCPep for predicting ncPEPs in various types of cancers in the 10 classes dataset.

|  | **without HyperGraph** | **with HyperGraph** |
| --- | --- | --- |
| *Anal canal cancer* | **0.5137** | **0.6619** |
| *Bladder cancer* | **0.5819** | **0.6980** |
| *Breast cancer* | **0.5891** | **0.7226** |
| *Colon cancer* | **0.5966** | **0.6333** |
| *Kidney cancer* | **0.6181** | **0.7430** |
| *Leukemia* | **0.5056** | **0.7344** |
| *Lung cancer* | **0.6028** | **0.6748** |
| *Prostate cancer* | **0.5606** | **0.6907** |
| *Skin cancer* | **0.5579** | **0.6411** |
| *Tongue cancer* | **0.6247** | **0.8317** |

**Table S3.** Performance evaluation on the MCC, ACC and AUC of HGCPep, CNN, GRU, LSTM, LSTM with Attention, RNN and CNN for predicting ncPEPs in the 15 classes dataset.

|  | **without HyperGraph** |  |  | **with HyperGraph** |  |  |
| --- | --- | --- | --- | --- | --- | --- |
|  | **MCC** | **ACC** | **AUC** | **MCC** | **ACC** | **AUC** |
| **CNN** | **0.0331** | **0.7640** | **0.5149** | **0.2770** | **0.8209** | **0.0331** |
| **GRU** | **0.0308** | **0.5464** | **0.5240** | **0.2795** | **0.7972** | **0.0308** |
| **LSTM** | **0.0304** | **0.5716** | **0.5239** | **0.2905** | **0.8096** | **0.0304** |
| **LSTM with**<br>**Attention** | **0.0933** | **0.7735** | **0.5449** | **0.2491** | **0.8063** | **0.0933** |
| **RNN and CNN** | **0.0059** | **0.7849** | **0.5025** | **0.3203** | **0.8071** | **0.0059** |
| **HGCPep**<br>**(Ours)** | **0.1841** | **0.7894** | **0.5881** | **0.3508** | **0.7720** | **0.7161** |

**Table S4.** Performance evaluation on the MCC, ACC and AUC of HGCPep, CNN, GRU, LSTM, LSTM with Attention, RNN and CNN for predicting ncPEPs in the 10 classes dataset.

|  | **without HyperGraph** |  |  | **with HyperGraph** |  |  |
| --- | --- | --- | --- | --- | --- | --- |
|  | **MCC** | **ACC** | **AUC** | **MCC** | **ACC** | **AUC** |
| **CNN** | **0.0295** | **0.6846** | **0.5147** | **0.3244** | **0.7779** | **0.6675** |
| **GRU** | **0.1005** | **0.6896** | **0.5504** | **0.2991** | **0.7575** | **0.6564** |
| **LSTM** | **0.0452** | **0.5291** | **0.5292** | **0.2695** | **0.7343** | **0.6471** |
| **LSTM with**<br>**Attention** | **0.1183** | **0.7244** | **0.5553** | **0.2356** | **0.7323** | **0.6230** |
| **RNN and CNN** | **-0.0062** | **0.7264** | **0.4988** | **0.3423** | **0.7609** | **0.6846** |
| **HGCPep**<br>**(Ours)** | **0.1506** | **0.7209** | **0.5751** | **0.3416** | **0.7107** | **0.7031** |

**Table S5.** Performance evaluation of the various hypergraph modules employed by HGCPep for forecasting ncPEPs in various cancers in the 15 classes dataset.

|  | **baseline**<br>**(without**<br>**HyperGraph)** | **with HGNN** | **with HGNNP** |
| --- | --- | --- | --- |
| ***Anal canal cancer*** | **0.5639** | **0.5666** | **0.5517** |
| ***Bile duct cancer*** | **0.5995** | **0.6639** | **0.6744** |
| ***Bladder cancer*** | **0.5806** | **0.7018** | **0.6695** |
| *Breast cancer* | **0.6293** | **0.6499** | **0.6452** |
| *Colon cancer* | **0.6066** | **0.6283** | **0.6579** |
| *Gastric cancer* | **0.6119** | **0.8254** | **0.8466** |
| *Kidney cancer* | **0.6387** | **0.6638** | **0.7225** |
| *Leukemia* | **0.5382** | **0.6945** | **0.7054** |
| *Liver cancer* | **0.5764** | **0.6606** | **0.7027** |
| *Lung cancer* | **0.5902** | **0.6644** | **0.6653** |
| *Ovary cancer* | **0.5277** | **0.7536** | **0.8142** |
| *Prostate cancer* | **0.5360** | **0.6671** | **0.6586** |
| *Skin cancer* | **0.6311** | **0.7146** | **0.6927** |
| *Thyroid cancer* | **0.5119** | **0.8622** | **0.8911** |
| *Tongue cancer* | **0.6793** | **0.8379** | **0.8434** |

**Table S6.** Performance evaluation of the various hypergraph modules employed by HGCPep for forecasting ncPEPs in various cancers in the 10 classes dataset.

|  | **baseline**<br>**(without**<br>**HyperGraph)** | **with HGNN** | **with HGNNP** |
| --- | --- | --- | --- |
| *Anal canal cancer* | **0.5137** | **0.6186** | **0.6619** |
| *Bladder cancer* | **0.5819** | **0.7256** | **0.6980** |
| *Breast cancer* | **0.5891** | **0.7004** | **0.7226** |
| *Colon cancer* | **0.5966** | **0.6483** | **0.6333** |
| *Kidney cancer* | **0.6181** | **0.7331** | **0.7430** |
| *Leukemia* | **0.5056** | **0.7309** | **0.7344** |
| *Lung cancer* | **0.6028** | **0.6642** | **0.6748** |
| *Prostate cancer* | **0.5606** | **0.6806** | **0.6907** |
| *Skin cancer* | **0.5579** | **0.7264** | **0.6411** |
| *Tongue cancer* | **0.6247** | **0.8457** | **0.8317** |

**Table S7.**
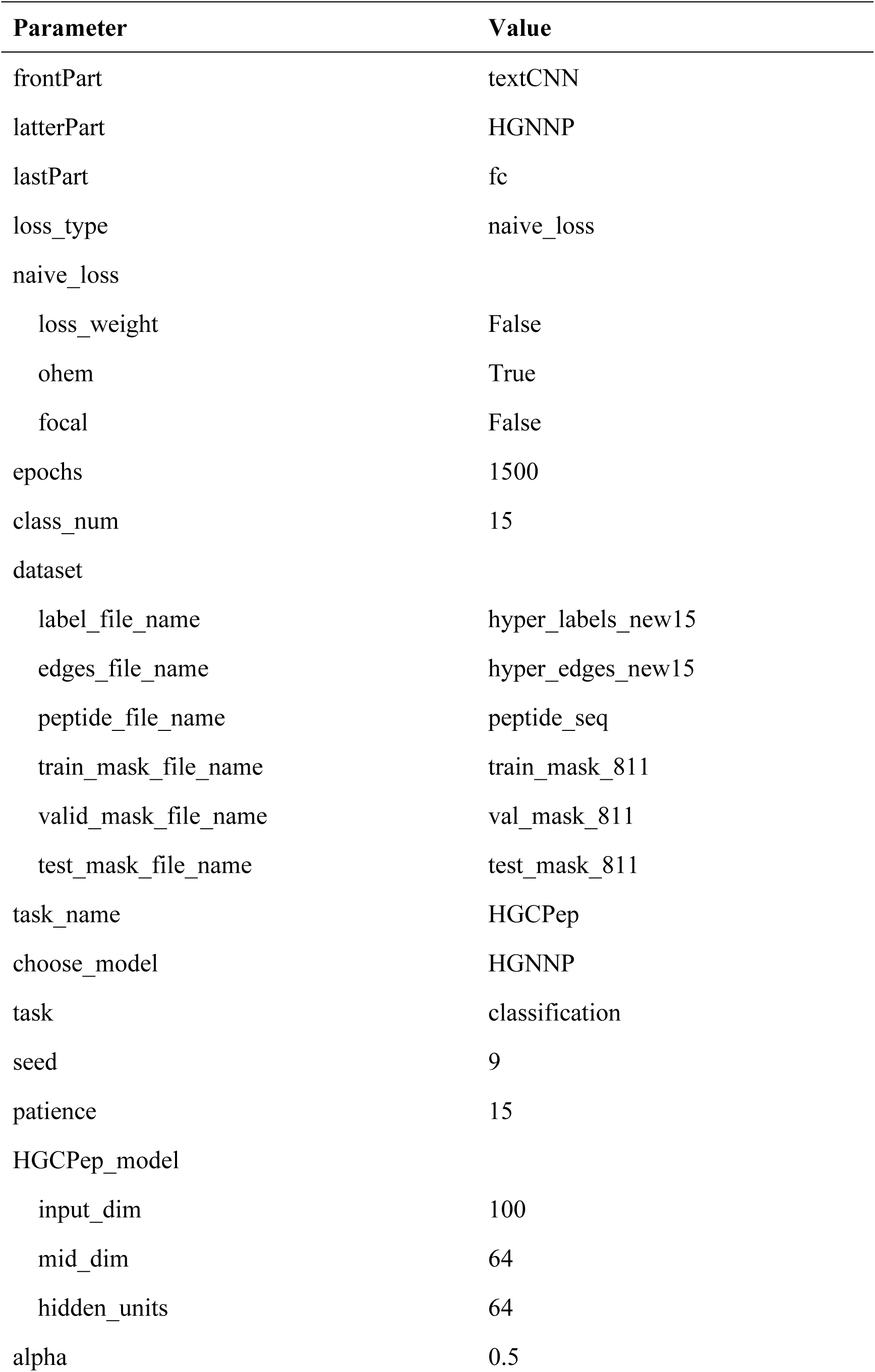
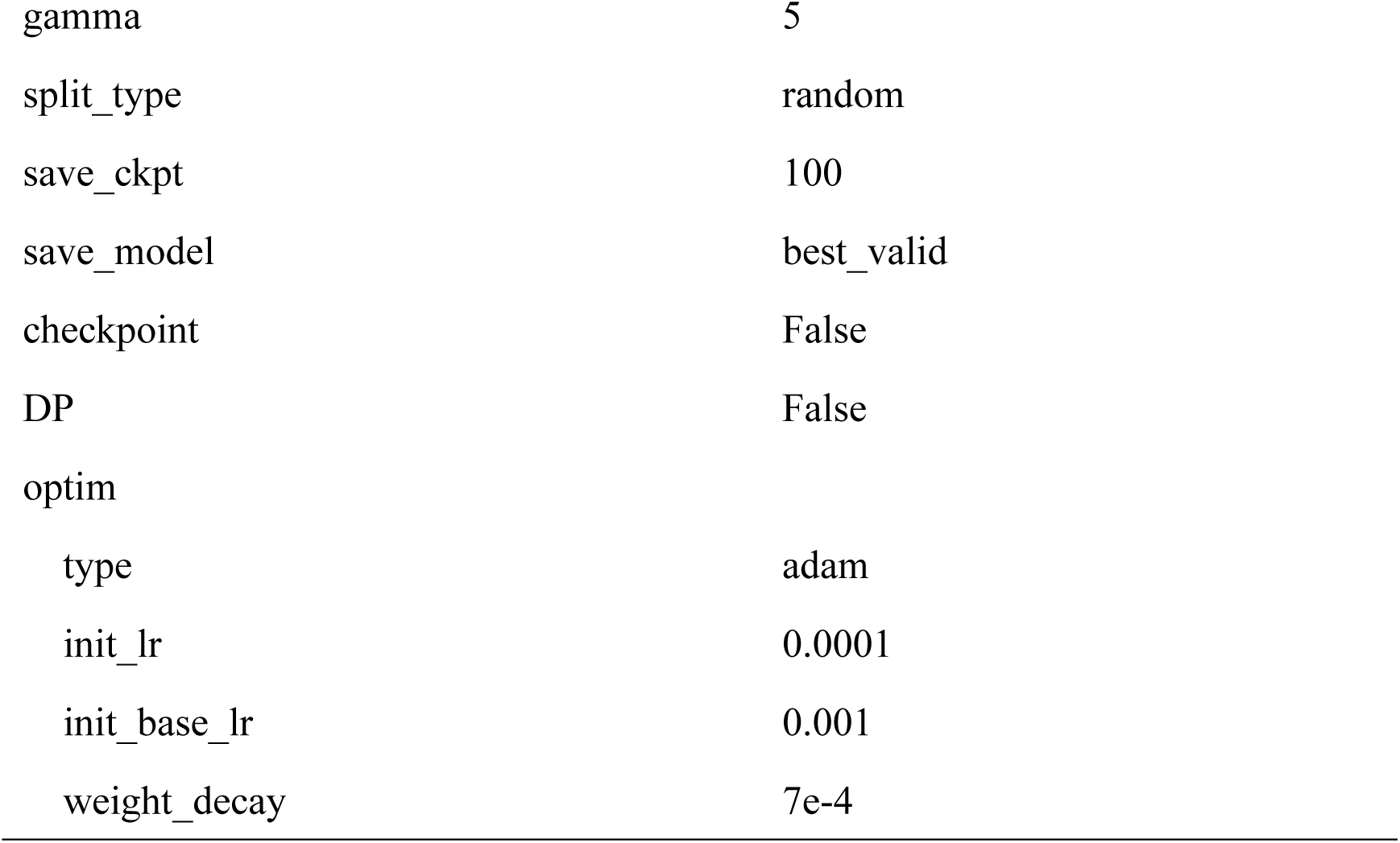
Experimental setup.

**Table S8.** Performance evaluation on the PREC and REC of HGCPep, CNN, GRU, LSTM, LSTM with Attention, RNN and CNN for predicting ncPEPs in the 15 classes dataset.

|  | without HyperGraph |  | with HyperGraph |  |
| --- | --- | --- | --- | --- |
|  | PREC | REC | PREC | REC |
| **CNN** | **0.2138** | **0.1734** | **0.4654** | **0.3820** |
| **GRU** | **0.1968** | **0.5420** | **0.4019** | **0.4620** |
| **LSTM** | **0.1944** | **0.4911** | **0.4342** | **0.4321** |
| **LSTM with Attention** | **0.2625** | **0.2191** | **0.4174** | **0.3709** |
| **RNN and CNN** | **0.1716** | **0.1039** | **0.4218** | **0.4944** |
| **HGCPep (Ours)** | **0.3419** | **0.3235** | **0.3893** | **0.6561** |

**Table S9.**
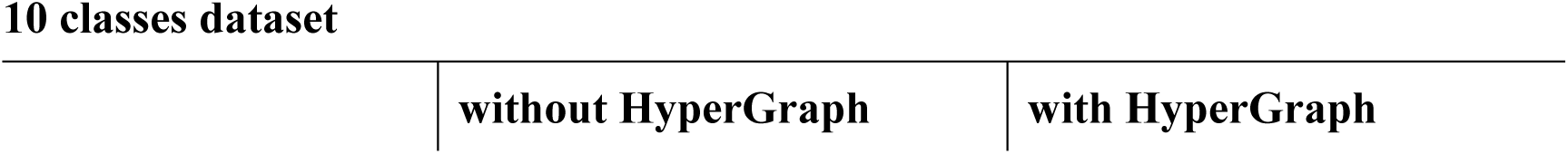
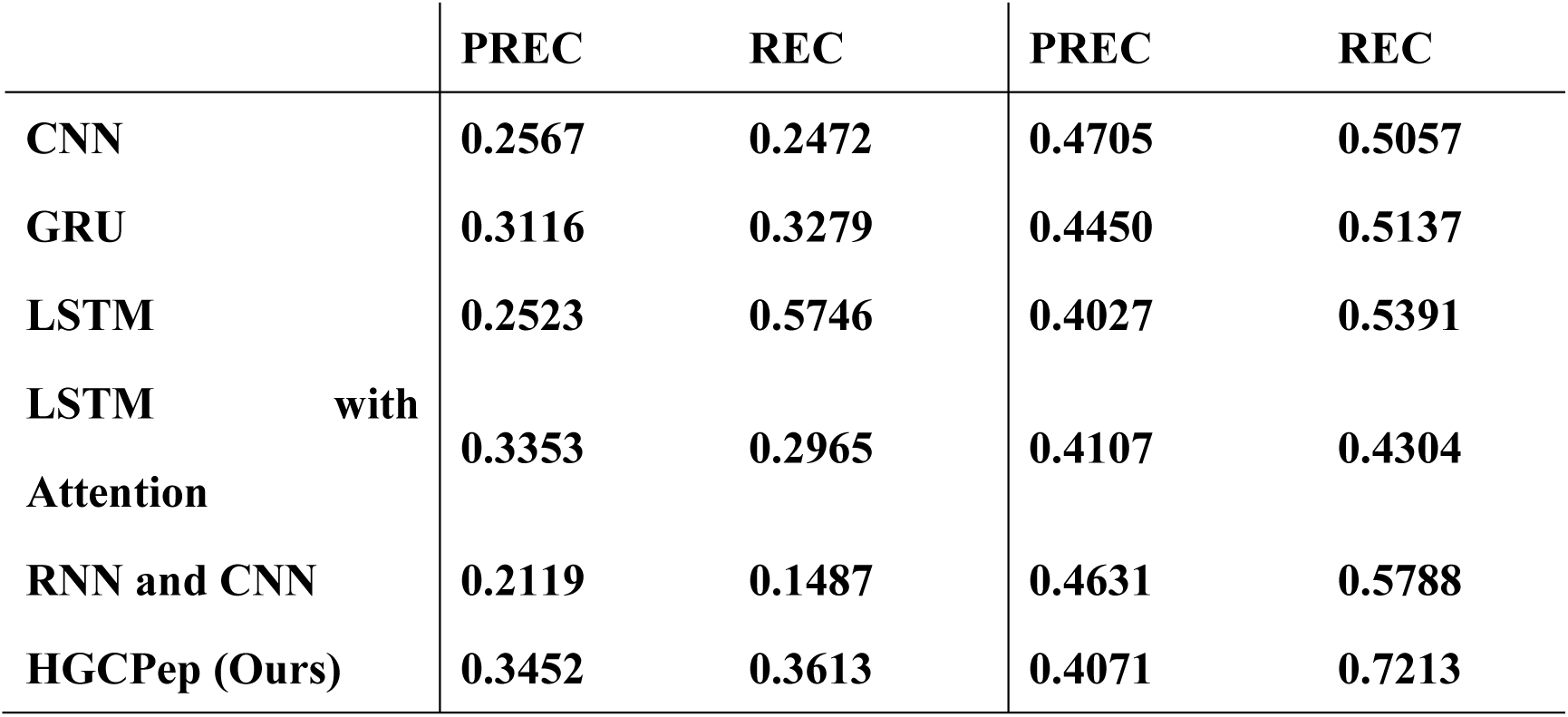
Performance evaluation on the PREC and REC of HGCPep, CNN, GRU, LSTM, LSTM with Attention, RNN and CNN for predicting ncPEPs in the 10 classes dataset.

|  | without HyperGraph | with HyperGraph |
| --- | --- | --- |

|  | PREC | REC | PREC | REC |
| --- | --- | --- | --- | --- |
| CNN | 0.2567 | 0.2472 | 0.4705 | 0.5057 |
| GRU | 0.3116 | 0.3279 | 0.4450 | 0.5137 |
| LSTM | 0.2523 | 0.5746 | 0.4027 | 0.5391 |
| LSTM with Attention | 0.3353 | 0.2965 | 0.4107 | 0.4304 |
| RNN and CNN | 0.2119 | 0.1487 | 0.4631 | 0.5788 |
| HGCPep (Ours) | 0.3452 | 0.3613 | 0.4071 | 0.7213 |

